# Olive mill solid waste induces beneficial mushroom-specialized metabolite diversity revealed by computational metabolomics strategies

**DOI:** 10.1101/2024.02.09.579616

**Authors:** Soliman Khatib, Idan Pereman, Elizabeth Kostanda, Mitja M. Zdouc, Nirit Ezov, Ron Schweitzer, Justin J. J. van der Hooft

## Abstract

**Introduction:** Mushrooms contain besides proteins a diverse pallet of specialized metabolites bioactive in either beneficial or harmful manner. Therefore, mushrooms have been exploited by humans for centuries for dietary or medical purposes. For example, the edible and medicinal mushrooms *Hericium erinaceus* and *Pleurotus eryngii* are grown commercially around the world. In nature, *H. erinaceus* grows on old or dead tree trunks, and *P. eryngii* grows on Apiaceae plant roots, whereas in cultivation, they grow on substrates mainly consisting of dry wood chips, straw, and cereals. To make their farming more sustainable, supplements such as olive mill solid waste (OMSW) have been added to support mushroom development. However, so far, the impact of substrate additives on the edible mushroom metabolic content has not been assessed.

**Methods:** Here, we examined the effect of different proportions of OMSW added to the substrate on the metabolic profiles of the fruiting body (FB) and mycelium of *H. erinaceus* and *P. eryngii* mushrooms. We used computational metabolomics strategies including GNPS molecular networking, MS2Query, and the FERMO dashboard, to organize, annotate, and prioritize metabolite features from the untargeted Q-Exactive Plus HR-LC-MS/MS metabolomics data. Following chromatography-based fractionation, the metabolite annotation of four metabolite features was further validated or fine-tuned using ^1^H-NMR, to resolve structural isomers.

**Results & Discussion:** Our computational metabolomics strategies showed several annotated metabolite features to be affected by OSMW concentration. In general, the methanolic extracts of *H. erinaceus* FB and mycelium were more highly enriched with specialized metabolites than those of *P. eryngii*. Interestingly, OMSW increased several hericenone analogues in the *H. erinaceus* FB, to which beneficial properties, such as anti-inflammatory, anticancer and neuroprotective properties are assigned, as well as several erinacerin metabolites from the mycelium. In addition, high concentrations of OMSW decreased the toxic enniatin metabolite abundance. In conclusion, we demonstrate how a change in substrate composition affects the mushroom’s specialized metabolome and can induce beneficial mushroom metabolite diversity. These results highlight the importance of including computational metabolomic strategies to investigate new sustainable growth options for edible mushrooms and other natural foods.

## Introduction

Mushrooms are popular food ingredients, not only due to their unique flavor and texture, but also due to their nutritious protein content and beneficial effects on human health^1,2^. For thousands of years, mushrooms have been used to support health, especially in East Asian countries^3^. In recent years, main biological constituents in edible and medicinal mushrooms have been explored for their health-related beneficial properties, both macromolecules, e.g., polysaccharides, polysaccharide–proteins/peptides and proteins, as well as low-molecular-weight molecules, e.g., cerebrosides, isoflavones, catechols, amines, triacylglycerols, sesquiterpenes and steroids^4,5^.

An example of an edible and medicinal mushroom is *Hericium erinaceus* (*H. erinaceus*) that is widely consumed in Asian countries^6^. Its fruiting body (FB) and mycelium are used in traditional Chinese medicine for the treatment of gastritis and hyperglycemia^7^. The pharmacological benefits of *H. erinaceus*, including antiaging, antioxidant, antitumor, antidiabetic, antidementia, antidepression^8^ and antianxiety activities^9–12^, are due to its large number of bioactive specialized metabolites, such as phenols (i.e., hericenones), aromatic compounds (i.e., hericerins, erinacerins, and erinaceolactones), sterols, polysaccharides and glycoproteins^13,14^. Another edible mushroom is *Pleurotus eryngii* (*P. eryngii*) that is cultivated for commercial use in many countries around the world. It has rapidly become a highly valued species in North Africa, Europe and Asia, in part due to its meaty structure that makes it a popular meat substitute. Many extracts from *P. eryngii* have been studied *in vivo* and *in vitro*, demonstrating interesting biological activities of the mushroom, including anticancer, antioxidant, antimicrobial, hypoglycemic and immunostimulatory effects^15–17^.

In nature*, H. erinaceus* grows on old or dead trunks of hardwood trees. *P. eryngii* grows on the roots of plants belonging to the *Apiaceae* family. Typically, in an industrial setting, mushrooms are grow indoors on substrates that mainly include dry plant materials (sawdust, dry wood chips, cereals, etc.), which commonly originate from agricultural and industrial wastes. Usually, a nitrogen source such as soybean hulls or grains can be added to the substrate to promote optimal mushroom development. Unfortunately, the olive oil industry produces extensive amounts of highly polluting by-products, both water waste and solid waste. Olive oil extraction can generate up to 30–40% Olive Mill Solid Waste (OMSW) and much effort has been invested in its exploitation^20,21^.To find a way to reduce the ecological impact of both mushroom cultivation and polluting waste products originating from the olive oil industry, OMSW was tested as an efficient nitrogen-contributing component^18,19^. Here, we do note that the type or characteristics of the growing substrate significantly influence the content of certain bioactive compounds in the mushrooms^22,23^. For example, recent studies have exhibited an increase in the healthy and beneficial glucans alpha and beta concentrations in correlation with the addition of several concentrations of OMSW to the substrate when growing *P. eryngii*^24^. However, a comprehensive assessment of the impact of OMSW on the mushroom’s specialized metabolome has not been performed up to date. Hence, in the present study, we grew *H. erinaceus* and *P. eryngii* on a mushroom substrate mixed with OMSW to explore the effect of OMSW addition on the mushroom’s metabolite profile and the possible impact on the consumers that may have.

Untargeted metabolomics based high-resolution (HR) LC-MS/MS is a widely-used method for analyzing small-molecule metabolites present in a biological sample in a particular physiological state, currently one of the most rapidly evolving research fields. Untargeted metabolomics can provide a “snapshot” of the sample, by identifying and quantifying small molecules metabolites. Many studies have made use of untargeted metabolomics to analyze plant samples, and some have investigated mushrooms^25,26^. To facilitate the analysis and interpretation of the complex datasets obtained from HRLC-MS/MS in untargeted metabolomics, computational metabolomics strategies are employed. The area of computational metabolomics encompasses computational, statistical, and machine-learning methods to analyze and interpret metabolomic data and its integration with other datasets, such as various omics or clinical data^27,28^. In the present study, we assessed the effect of OMSW on metabolite-profile diversity in the FB and mycelium of *H. erinaceus* and *P. eryngii* mushrooms using computational metabolomics methods to analyze the HR LC-MS/MS data. Several in-silico metabolite annotations were further verified with 1D-1H-NMR of chromatographic fractions. The study results can be used to assess the impact of using various mushroom growth resources on their edible constituents. Moreover, the strategies employed here could be applied to other types of food and substrates as well.

## Methods

### Mushroom growth conditions

*P. eryngii* and *H. erinaceus* were grown on a sterilized mixture of eucalyptus sawdust (originating from forest management cut-wood piles in the upper Galilee in northern Israel) and OMSW (collected from a local olive-oil production factory), with the addition of malt waste (collected from a local Israeli beer production factory). OMSW, the solid fraction from a three-phase olive mill, was added at different concentrations of up to 80% (w/w) (Table 1). Also prepared was a control without OMSW (0% (w/w)). The mixture was wetted to 52–58% water content and packed into 2 L polypropylene bags containing a microporous filter (Unicornbags, USA type A4), resulting in 0.8 kg wet substrate/bag. The bags were autoclaved at 121°C for 1 h and cooled to 25°C for inoculation with grain spawn. The culture was incubated at 23.5°C for 14–21 days. For fruiting, the bags were opened, and the temperature was reduced to 18°C, with a relative humidity of 90% and 8 h of light daily.

**Table 1.** The four experimental groups of growth media with varying concentrations of Olive Mill Solid Waste (OMSW) with their constituents, as well as their estimated moisture (%).

|  | **OMSW 80%** | **OMSW 60%** | **OMSW 33%** | **OMSW 0%** |
| --- | --- | --- | --- | --- |
| OMSW (%) | 80 | 60 | 33 | 0 |
| Malt (%) | 10 | 20 | 33 | 50 |
| Eucalyptus (%) | 10 | 20 | 33 | 50 |
| Moisture<br>(Estimated %) | 52 | 54 | 58 | 60 |

FB were freshly harvested at the Matityahu Experimental Farm (Upper Galilee, Israel). They were collected according to their maturity level (spines elongation in *H. erinaceus*, cap opening in *P. eryngii*). After harvest, FB and spent mushroom substrate (SMS) were weighed, chopped into small pieces, and freeze-dried.

### Extraction conditions

All samples of FB, SMS, and mushroom substrate (MS) were ground using a mechanical grinder, briefly vortexed and extracted in methanol (MeOH; analytical, suitable for LC-MS) at a ratio of 1:20 (1 g of dried material in 20 ml MeOH). The extraction was performed by shaking at 25°C and 1000 rpm for 2 h. After extraction, the samples were centrifuged (4°C, 3220*g* for 30 min) and the liquid was collected and filtered through a 0.45-µm syringe filter; 1 mL of the filtered extract was kept in a LC-MS vial at - 80°C while the remaining extract was dried in an evaporator (for yield calculation). Dry extracts were kept at -20°C.

### HPLC analysis

The samples were analyzed by injecting 5 μl of the extracted solutions into a Dionex Ultimate 3000 ultra-HPLC system connected to a photodiode array detector (Thermo Fisher Scientific), with a reverse-phase column (ZORBAX Eclipse Plus C18, 100 x 3.0 mm, 1.8 μm; Agilent). The mobile phase consisted of (A) 0.1% formic acid in double distilled water (DDW) and (B) 0.1% formic acid in MeOH. The gradient began at 95% A which was kept isocratic for 1 min, then decreased to 40% B over 2 min and kept isocratic for 2 min, then increased to 97% B over 8 min and kept isocratic at 97% B for another 19.5 min. Phase A was returned to 95% over 1.5 min and the column was allowed to equilibrate at 95% A for 3 min before the next injection. The flow rate was 0.4 mL/min.

### LC–MS/MS analysis

MS/MS analysis was performed with a heated electrospray ionization (HESI-II) source connected to a Q Exactive Plus Hybrid Quadrupole-Orbitrap mass spectrometer (Thermo Fisher Scientific). ESI capillary voltage was set to 3000 V, capillary temperature to 350°C, gas temperature to 350°C and gas flow to 35 mL/min. The mass spectra (m/z 100–1000) were acquired in negative- and positive-ion modes with high resolution (FWHM = 70,000). MS^1^ parameters were: resolution 70,000, AGC target 3E^6^, maximum IT 100 ms, scan range 67–1000 m/z; MS^2^ parameters were: resolution 17,500, AGC target 1E^5^, maximum IT 50 ms, loop count 5, MSX count 1, isolated window 1, collision energy (N)CE 20, 50 and 80 EV; data dependent (dd) settings were: minimum AGC 8E^3^, apex trigger 6 to 16 s, exclude isotope on, dynamic exclude 8.0 s.

### Chromatographic fractionation and NMR Analysis

H. erinaceus FB (2g) grown with 80% OMSW were dried, ground, and extracted with 40 ml methanol for 2 h at room temperature to obtain 960 mg of crude extract. The crude extract was separated into fractions using a silica gel column (FP ECOFLEX Si 25g) connected to a pure chromatographic system, C-815 flash connected to UV and ELSD detectors from BUCHI-Switzerl. The mobile phase was a gradient of hexane-ethyl acetate and then ethyl acetate-methanol. Three fractions, fraction 1 (8.5 mg), fraction 2 (7.2 mg), and fraction 3 (4 mg), were dissolved in 1 ml deuterated chloroform (CDCl3) and 1H-NMR spectra were recorded at room temperature with a Bruker 400 MHz instrument with chemical shifts reported in ppm relative to the residual deuterated solvent. Transmitter freq.: 400.402472 MHz, time domain size: 65536 points, width: 8196.72 Hz = 20.4712 ppm = 0.125072 Hz/pt, number of scans: 16.

### Computational metabolomics analysis

#### Data reprocessing

Data-dependent MS/MS spectrum files were converted from .raw to .mzXML using the GNPS vendor-conversion (Mass Spectrometry File Conversion – GNPS Documentation; ccms-ucsd.github.io)^29^. Data preprocessing was performed using MZmine 3.3.0 software and included the following steps: mass1 and mass2 detection with noise levels of 4.0E^5^ and 1.0E^4^ respectively, LC-MS chromatogram building and resolution with intensity threshold of 9.0E^5^ and minimum highest intensity of 2.0E^6^, isotope grouping, alignment, isotope pattern filtering, gap filling and filtering out of duplicate peaks^30^, M/Z and RT tolerance were set to 5 and 0.5 respectively. More details about MZmine 3 parameters can be found in the batch file in the supplementary data. The aligned feature lists were exported to the GNPS platform as MS/MS files (.mgf format) and quantification tables (.csv format of aligned features and related chromatographic peak areas).

#### FERMO analysis

FERMO_0.8.8 dashboard^31^ (permanently available from https://zenodo.org/records/7565701) was used for the prioritization of molecular features from the MS data which were upregulated or downregulated upon addition of OMSW. The MS/MS file (.mgf format) and full quantification table (.csv format) were obtained from MZmine 3 analysis using the ’Export molecular networking files (e.g., GNPS, FBMN, IIMN, MetGem)’ export button. Feature intensity was set to ‘Height’. The parameters used for the FERMO analysis were: mass deviation 5; minimum number of fragments per MS² spectrum 5; QuantData factor 2; Blank factor 10; relative intensity filter range 0.01–0.95; spectral similarity networking algorithm, modified-cosine; fragment similarity tolerance 0.1; fragment similarity tolerance 0.8. MS2Query was used to annotate the MS/MS data^32^.

#### Molecular networking analyses

Molecular networks were obtained using the feature-based molecular networking (FBMN) workflow^33^ in GNPS (https://gnps.ucsd.edu,). The precursor ion mass tolerance was set to 0.02 Da and the MS/MS fragment ion tolerance to 0.01 Da. A molecular network was then created where edges were filtered to have a cosine score above 0.6 and more than 4 matched peaks. The maximum size of a molecular family was set to 100. The spectra in the network were then searched against GNPS spectral libraries^29,34^. The library spectra were filtered in the same manner as the input data. All retained matches between network spectra and library spectra were required to have a score above 0.5 and at least 4 matched peaks. DEREPLICATOR+^35^, Network Annotation Propagation (NAP)^36^ and Structural Similarity Network Annotation Platform for Mass Spectrometry (SNAP-MS) were used to annotate MS/MS spectra^37^. The molecular networks were visualized using Cytoscape software^38^.

#### MolNetEnhancer Workflow for chemical class annotation of molecular networks

To elaborate on the chemical structure information in a molecular network, information from in-silico structure annotations generated by GNPS library search, NAP and DEREPLICATOR+ was incorporated into the network using the GNPS MolNetEnhancer workflow^39^ (https://ccms-ucsd.github.io/GNPSDocumentation/molnetenhancer/) on the GNPS website (http://gnps.ucsd.edu). Chemical class was annotated using ClassyFire chemical ontology^40^.

### Data deposition and job accessibility

The MS data were deposited in the public repository MassIVE: positive ionization mode in (<u>MSV000090919</u>), and negative ionization mode in (<u>MSV000090920</u>). The molecular networking jobs can be publicly accessed and browsed at the following URLs: https://gnps.ucsd.edu/ProteoSAFe/status.jsp?task=069e4c5dc7d846a4a96e485091d199a4 for *H. erinaceus*, https://gnps.ucsd.edu/ProteoSAFe/status.jsp?task=60727fe5228643e6a482bd797d83df38 for *P. eryngii,* and https://gnps.ucsd.edu/ProteoSAFe/status.jsp?task=2b86dd35cc4a4219bad07c3519ad78bf for *H. erinaceus* and *P. eryngii* together.

## Results and discussion

*H. erinaceus* mushrooms were grown on sterilized substrate comprised of eucalyptus sawdust and malt waste mixed with different percentages of OMSW (0%, 33%, 60% and 80%). FB, SMS and mushroom substrates were collected, extracted, and analyzed by HRLC-MS/MS. Mushroom substrates with 0%, 33%, 60% and 80% OMSW were used as blanks to filter out non-biological signals. The effect of OMSW on mushroom growth and development was reported previously ^18,19^. In the present study, the impact of the OMSW on mushrooms specialized metabolite diversity was examined using computational metabolomics approaches.

The use of MZmine 3 for LC-MS/MS data processing in positive ionization mode led to the detection of 2766 and 1899 features in the *H. erinaceus* and *P. eryngii* mushroom sample extracts, respectively, for which MS/MS data was collected and that were further analyzed by FBMN and subsequently prioritized using the FERMO dashboard.

### FERMO platform to prioritize *H. erinaceus* FB metabolites affected by OMSW

The priotorization software FERMO was run and the FERMO dashboard was used to prioritize metabolites that were upregulated or downregulated with increasing OMSW percentage in the mushroom substrate. Molecular features that were also detected in the mushroom substrate blanks containing 0%, 33% 60% and 80% OMSW, yellow-colored peaks in Figure 1A were removed as they cannot be mushroom-generated, leaving 360 features in the *H. erinaceus* FB extracts (Fig. 1B). From these, 56 features were selected with over 2-fold higher quantity (signal intensity/height) in the mushrooms grown with 80% OMSW compared to those grown with 0% (Fig. 1C and Fig. S1). Most of these selected features were connected in two molecular families based on mass spectral similarity networking, and their quantities increased dose-dependently with increasing percentage of the OMSW (Fig. S1). This increase was significant for 18 of the metabolite features (Fig. S2).

**Fig. 1.**
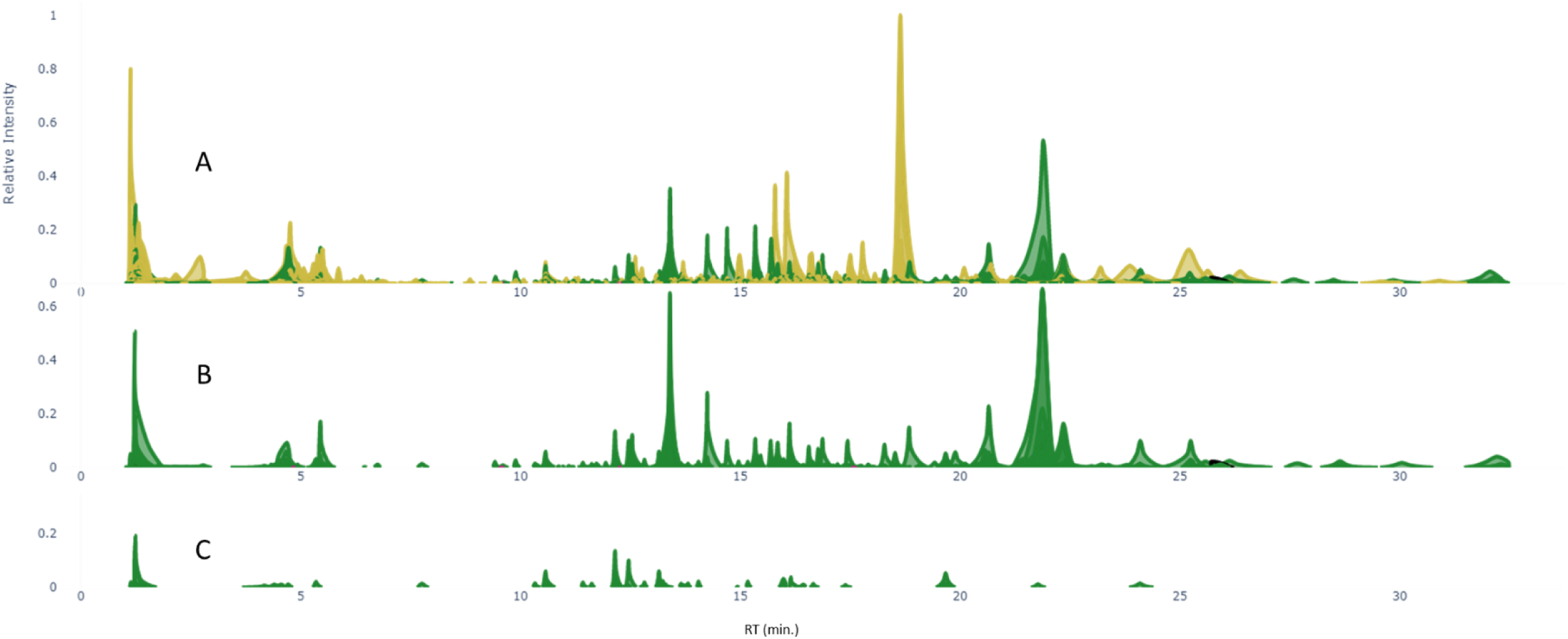
FERMO analysis of MS data from extracts of *H. erinaceus* mushrooms grown on substrate with different percentages of OMSW. (A) Pseudo chromatogram of molecular features detected in extracts of *H. erinaceus* fruiting bodies (FB) grown on mushroom substrate mixed with 80% OMSW (green), and features detected in the mushroom substrates medium blank samples (yellow). (B) Pseudo chromatogram of molecular features detected in extracts of *H. erinaceus* FB after blank removal. (C) Pseudo chromatogram of molecular features with 2-fold higher quantity in the FB extracts of *H. erinaceus* grown on 80% OMSW compared to 0% OMSW.

Three of the metabolites that increased with the addition of OMSW were annotated using MS2Query (Fig. 2 A-C), with a score higher than 0.7 and Δm/z = 0, as acetyl carnitine (Fig. 2A & 2G-1), (4aR,10aR)-6,7-dihydroxy-1,1,4a-trimethyl-3,4,10,10a-tetrahydro-2H-phenanthren-9-one (Fig. 2B & 2G-2) and 6-[(3E,5E,7S)-5,7-dimethyl-2-oxonona-3,5-dienyl]-2,4-dihydroxy-3-methylbenzaldehyde (Fig. 2C & G-3). Two other metabolites that did not significantly change upon the addition of OMSW (Fig. 2D-E) were annotated by DEREPLICATOR+ as the *Hericium*-unique metabolites hericenones F (Fig 2D & 2G-4) and hericenone H (Fig. 2E & 2G-5), with scores of 12 and 15, respectively. A metabolite whose quantity decreased significantly in a dose-dependent manner with the addition of OMSW was annotated by DEREPLICATOR+ to be pregn-5-ene-3,14,20-triol_3-O-[3-O-methyl-α-D-galactopyranosyl-(1→4)-β-D-digitoxopyranoside] (Fig. 2F & 2G-6) with a DEREPLICATOR+ score of 7. Interestingly, none of these metabolites were annotated with a MS2Query score higher than 0.7, showing the complementarity of existing annotation strategies. In addition, 64 metabolite features were detected mainly in the 0% OMSW group, with only traces of these metabolites detected in the FB grown on mushroom substrates mixed with OMSW (Fig. S3).

**Fig. 2.**
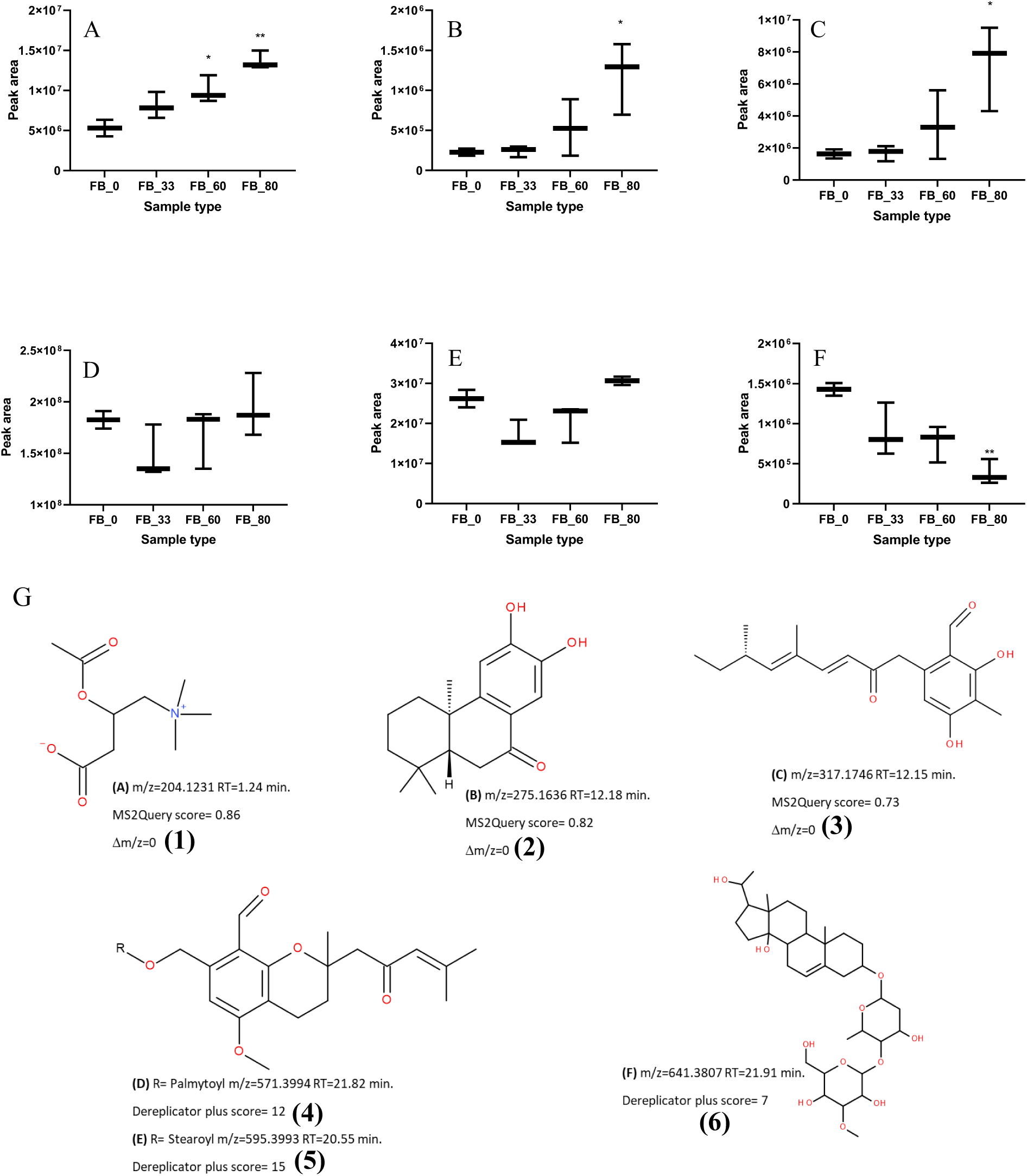
(A, B and C) Box and whisker plots of selected metabolites whose quantities were significantly increased in the *H. erinaceus* fruiting body (FB) with increasing percentage of OMSW in the mushroom substrate. (D and E) Metabolites whose quantities did not change in the FB with increasing OMSW in the mushroom substrate. (F) A metabolite that significantly decreased in the FB with increasing OMSW percentage in the mushroom substrate. The figures present the mean area values ± SD from three extracts for each group (*n* = 3). \**P* ≤ 0.05, \*\**P* ≤ 0.01 relative to the 0% OMSW FB group. Statistical analysis was carried out by one-way ANOVA and GraphPad Prism 9 software. (G) The metabolite structures were annotated using MS2Query or DEREPLICATOR+.

### FBMN analysis of *H. erinaceus* and *P. eryngii* mushroom extracts

Feature-based Molecular Networking (FBMN) was used to organize and annotate the LC-MS/MS data of the extracts obtained from the *H. erinaceus* and *P. eryngii* FB, SMS and mushroom substrates)without mycelium(. First, the LC-MS/MS data of the extracts was processed using MZmine 3 to obtain 2766 and 1899 features for *H. erinaceus and P. eryngii*, respectively followed by an analysis using FBMN on the GNPS platform. Of the obtained features, 1258 and 1837 were connected by molecular networks for *P. eryngii* and *H. erinaceus*, respectively (Figs. S4 A and B). After removing molecular networks that included features from mushroom substrates mixed with 0%, 33%, 60% and 80% OMSW without mushroom mycelium (blank features), 752 and 240 features remained for the FB and mycelium of *H. erinaceus* and *P. eryngii*, respectively (Fig. S4 C and D).

Of the 240 features obtained from the *P. eryngii* FBMN analysis, 153 were annotated using GNPS libraries, 33 of them with confidence higher than 95%; 153 were annotated using NAP and 142 were annotated using DEREPLICATOR+, 84 of the latter with a score higher than 10 (Fig. S5A). *P. eryngii* FB and mycelium exhibited a relatively limited number of secondary metabolites. Molecular family classification using ClassyFire showed that the majority belonged to phospholipids (22%), amino acid and carboxylic acid derivatives (15%), steroids (15%), sphingolipids (10%) and fatty acids (8%) (Fig. S5B).

Of the 752 features obtained from the *H. erinaceus* FBMN analysis, 165 were annotated using GNPS libraries, 28 of them with confidence higher than 95%; 165 were annotated using NAP and 406 were annotated using DEREPLICATOR+, 221 of the latter with a score higher than 10 (Fig. 3A). FBMN analysis showed that *H. erinaceus* FB and mycelium were richer in specialized secondary metabolites than *P. eryngii*. About 40% of the *H. erinaceus* secondary metabolites belonged to the prenol lipids, which distributed into coumarins, sesterterpenoids, triterpenoids, diterpenoids and diterpene lactones (Fig. 3 B and C).

**Fig. 3.**
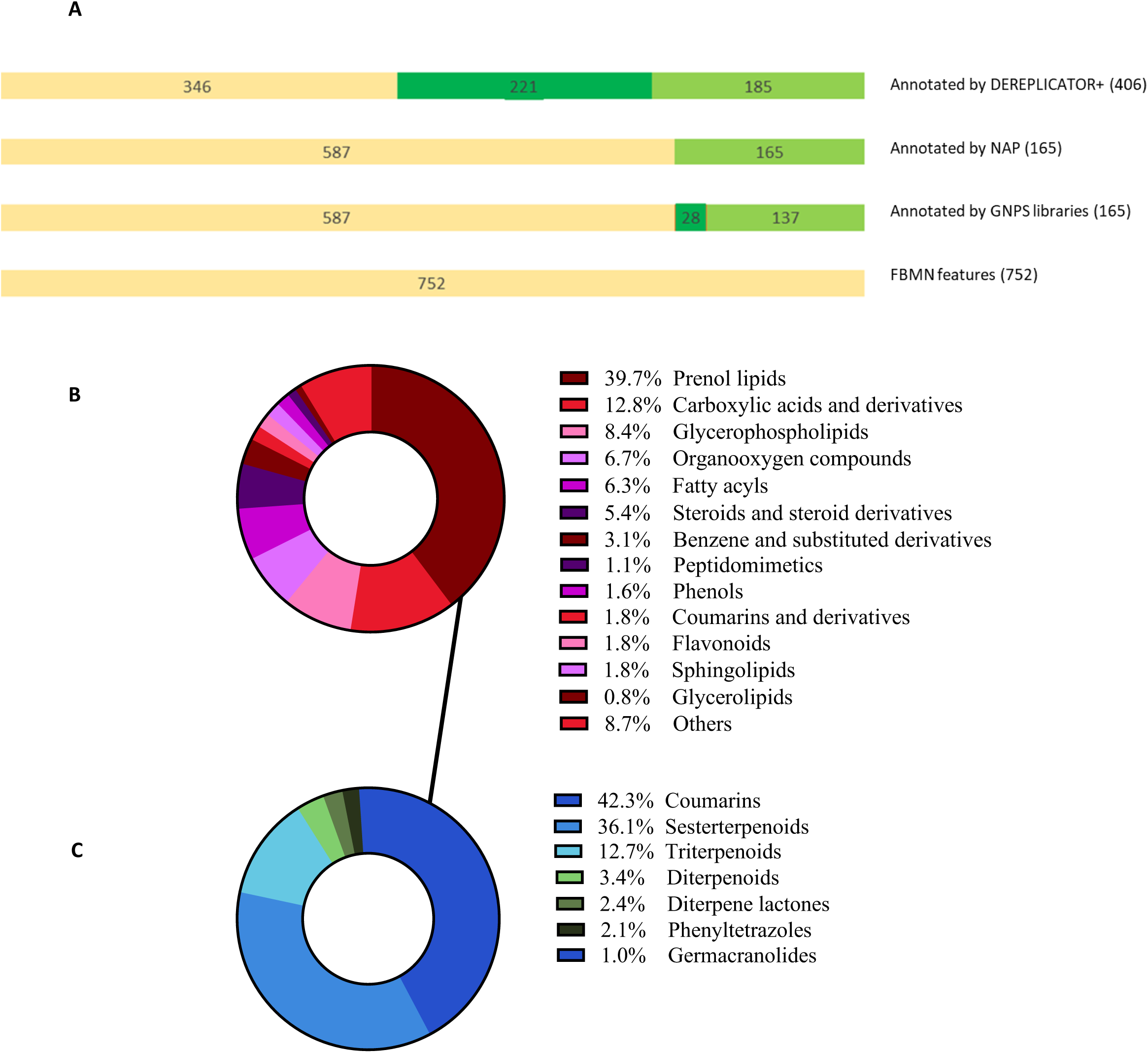
(A) Annotation results obtained from *H. erinaceus* FBMN analysis using GNPS libraries, DEREPLICATOR+ and NAP. The yellow color represents number of detected features, Green color the annotated features and dark green color the annotated features with score higher than 0.95 for GNPS libraries and higher than 12 for DEREPLECATOR plus database (B) Chemical classification (using ClassyFire) of the molecular families obtained from the *H. erinaceus* FBMN. (C) Molecular distribution of the prenol lipids obtained from *H. erinaceus* FBMN.

Several ergostane derivatives had been previously isolated from *H. erinaceus*, and some of their biological activities reported, such as anti-inflammatory effects and activation of the transcriptional activity of peroxisome proliferator-activated receptors (PPARs)^41^. Compounds that modulate the functions of PPARs are beneficial for the treatment of type 2 diabetes, obesity, metabolic syndromes, inflammation, and cardiovascular disease ^42^. In our study, one molecular family contained derivatives and analogs of interesting secondary metabolites belonging to the ergostane-type sterol fatty acid esters as annotated by both DEREPLICATOR+ and SNAP-MS. The nodes of the molecular family showed that these compounds are not specific to *H. erinaceus* but were detected in both *H. erinaceus* and *P. eryngii* in both the FB and mycelium (Fig. 4). In contrast to the ergostane family, most of the other specialized metabolites were unique to *H. erinaceus* FB or *H. erinaceus* mycelium.

**Fig. 4.**
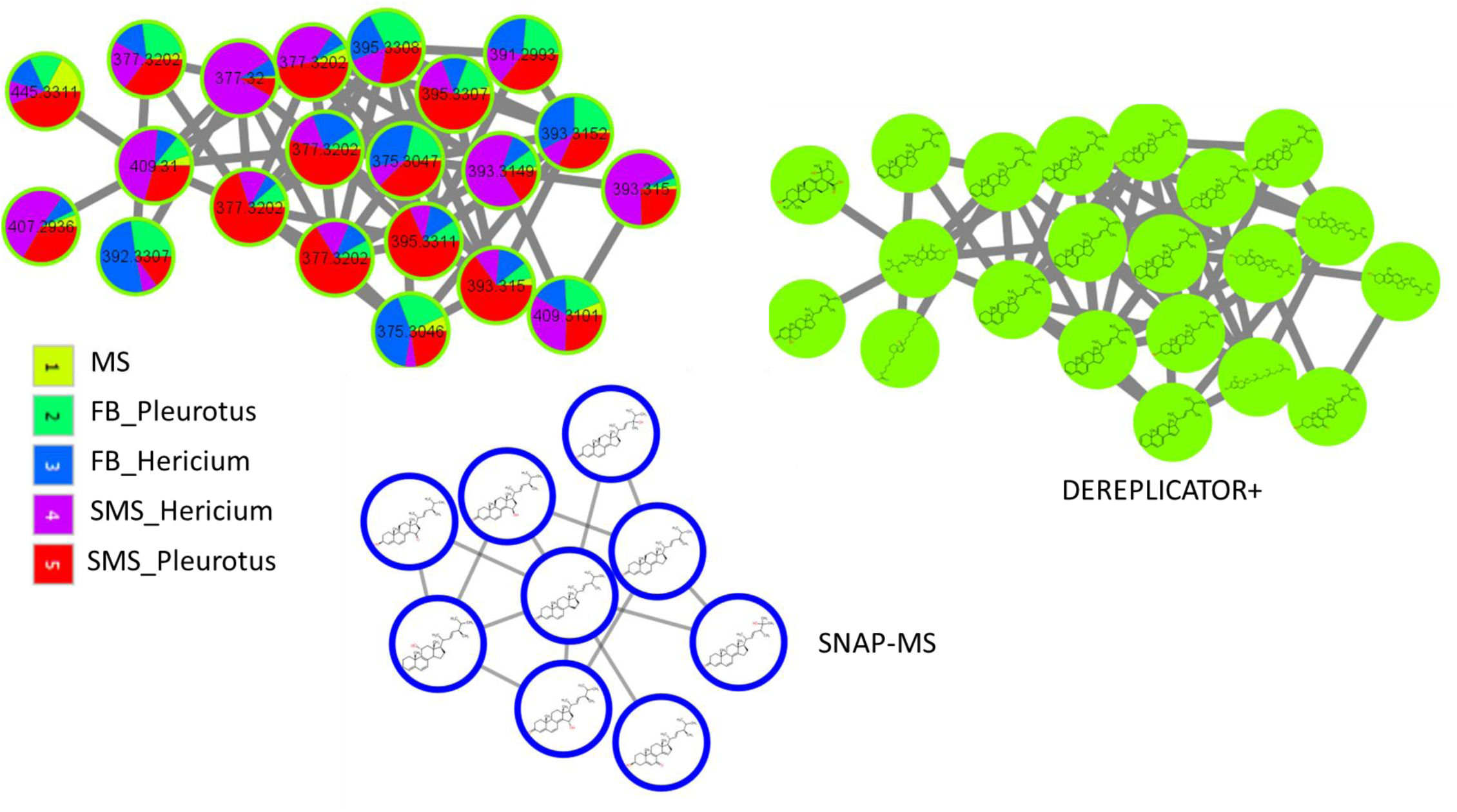
Molecular families from the FBMN results, with most components detected in both *H. erinaceus* and *P. eryngii* fruiting bodies (FB) and mycelium (SMS). Most of the components were annotated as ergosterol derivatives and analogs using both DEREPLICATOR+ and SNAP-MS such as, Ergosta-5,7,9(11),22-tetraen-3β-ol, Ergosta-3,5,7,9(11),22-pentaene and Citreoanthrasteroid_B.

It has been reported that more than 60 bioactive specialized metabolites are unique to *H. erinaceus* FB or mycelium with antitumor, antibacterial, hypoglycemic and neuroprotective effects^25,43,44^. These specialized secondary metabolites include structure families of erinacerins, erinacines, hericenones, hirecenes and ergostanes. In our study, none of these reported metabolites were annotated using GNPS libraries or NAP; however, of the bioactive metabolite groups mentioned above, the in-silico annotation tool DEREPLICATOR+ could annotate 14, and SNAP-MS could annotate 26 of them (Fig. 5).

**Fig. 5.**
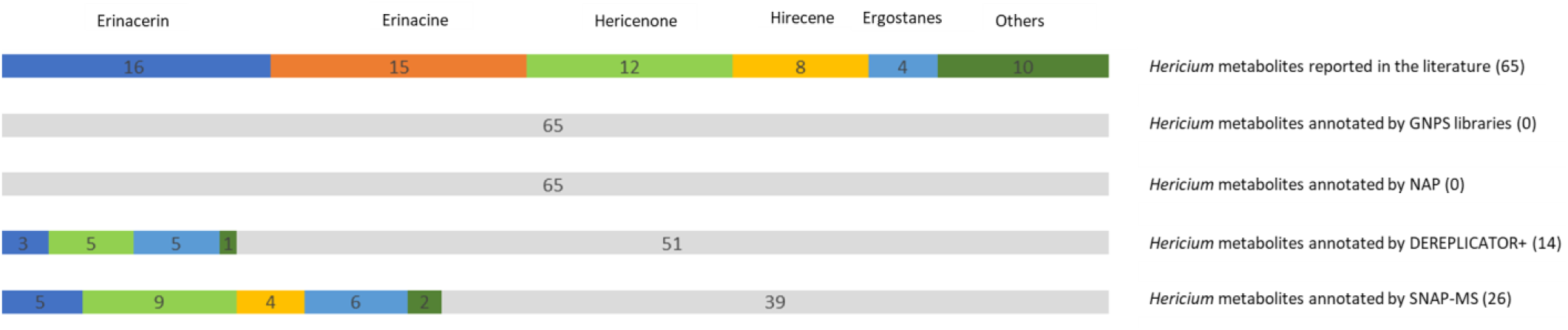
Unique metabolite families isolated from *H. erinaceus* and annotated using GNPS libraries, NAP, DEREPLICATOR+ and SNAP-MS.

Hericenones are specialized metabolites uniquely found in *H. erinaceus*^12,13^; more than 10 hericenone compounds (A–J) and isohericenone J have been isolated from the FB of *Hericium*: Hericenones C, D and E have been reported to have a stimulatory effect on the synthesis of nerve growth factor^45^, hericenone F to have anti-inflammatory activity^46^, and hericenone J and isohericenone J to have anticancer activities^47^.

In accordance with the literature, our FBMN results showed two molecular families that were unique to the *H. erinaceus* FB extracts^48^. Their components were classified to belong to hericenone derivatives and analogs as annotated by DEREPLICATOR+ and SNAP-MS. Five hericenones were annotated using DEREPLICATOR+ and nine using SNAP-MS. MS2LDA results showed that the nodes of the molecular family in Fig. 6A share the unsupervised Mass2Motifs 231 and 155 (tentative substructure annotations in Fig. S6). The molecular family in Fig. 6B contains hericenone J analogs (m/z = 317.1745), as annotated by SNAP-MS. We also note that in accordance with the FERMO results, most of the nodes of these two molecular families were upregulated by increasing the OMSW percentage in the mushroom substrate.

**Fig. 6.**
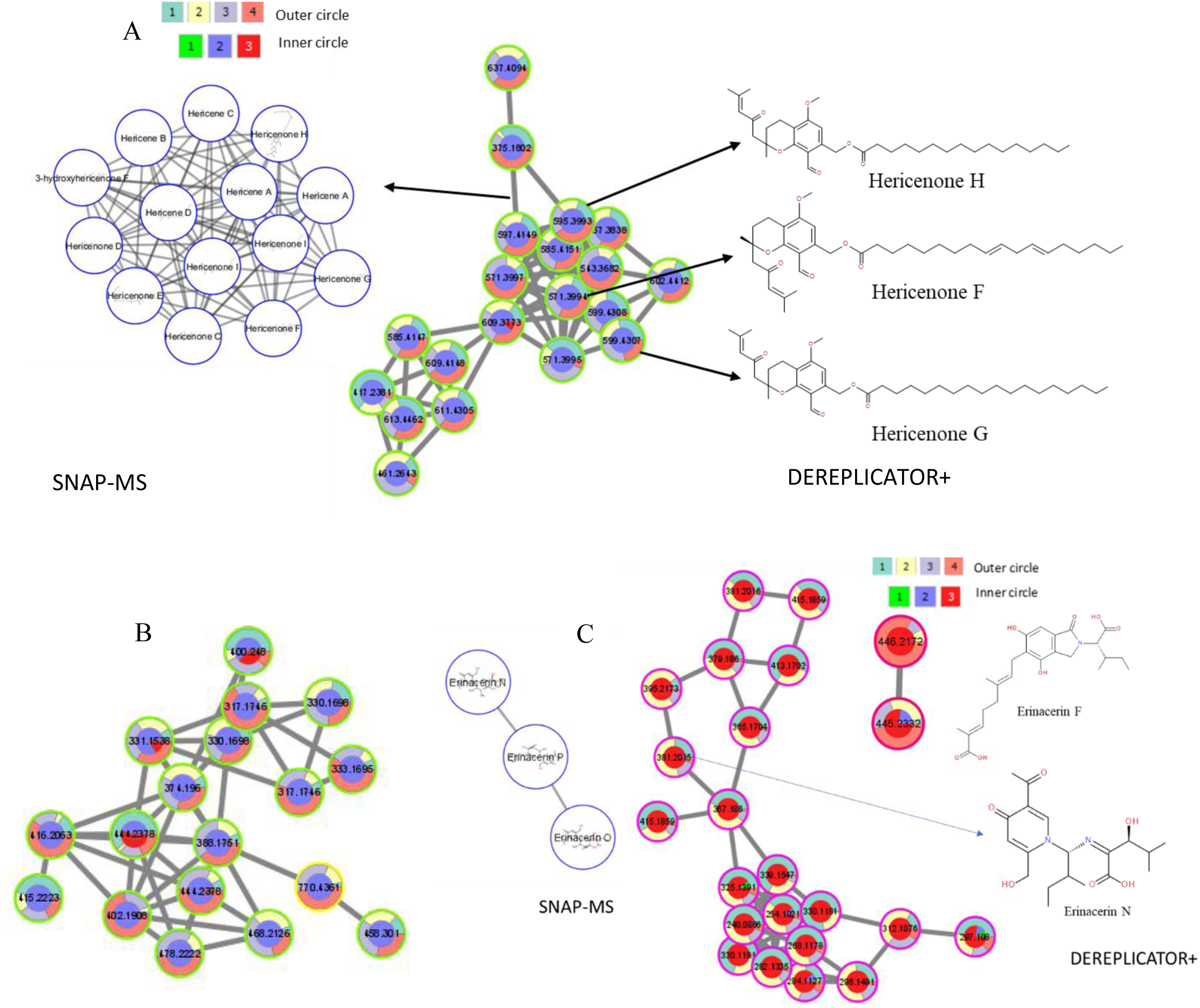
Cytoscape visualization of the molecular features analyzed by FBMN mass spectral similarity networking. Colors of the inner circle represent features detected in the mushroom substrate (MS), fruiting body (FB) and spent mushroom substrate (SMS) – 1, 2 and 3, respectively. Colors of the outer circle represent features detected in the groups of 0, 33, 60 and 80% OMSW – 1, 2, 3 and 4, respectively. (A and B) Molecular families whose components were detected mostly in the FB and increased with increasing OMSW percentage. (C) Molecular families whose components were detected mostly in the mycelium, some of them increasing with the addition of OMSW.

Furthermore, Erinacerins are specialized metabolites with a unique mode of action as they inhibit α-glucosidase, a target used to treat diabetes type II^49^. More than 15 erinacerins (A-O) were isolated from the *H. erinaceus* mycelium. In accordance with the literature, our FBMN results complemented with DEREPLICATOR+ and SNAP-MS annotations showed that most of the annotated erinacerins and their analogs were isolated from the SMS that contains the mycelium^50,51^. In addition, some of the molecular family nodes related to erinacerin analogs had increased abundance with increasing OMSW percentage in the mushroom substrate (Fig. 6C).

Another two molecular families detected in the *H. erinaceus* FB extracts and the nodes of one of the molecular families were classified by SNAP-MS as belonging to enniatin derivatives and analogs (Fig. 7A). Enniatins are toxic due to their ability to act as ionophores, changing ion transport across membranes and disrupting the ionic selectivity of cell walls. In the membrane, enniatins form a dimeric structure which transports ions (especially K^+^, Mg^2+^, Ca^2+^ and Na^+^) across membranes. The other molecular family components were not annotated (Fig. 7B). In accordance with the FERMO results, the compounds related to these two molecular families were downregulated by the addition of OMSW, and most of their metabolites only appeared in the 0% OMSW group (Fig. 7).

**Fig. 7.**
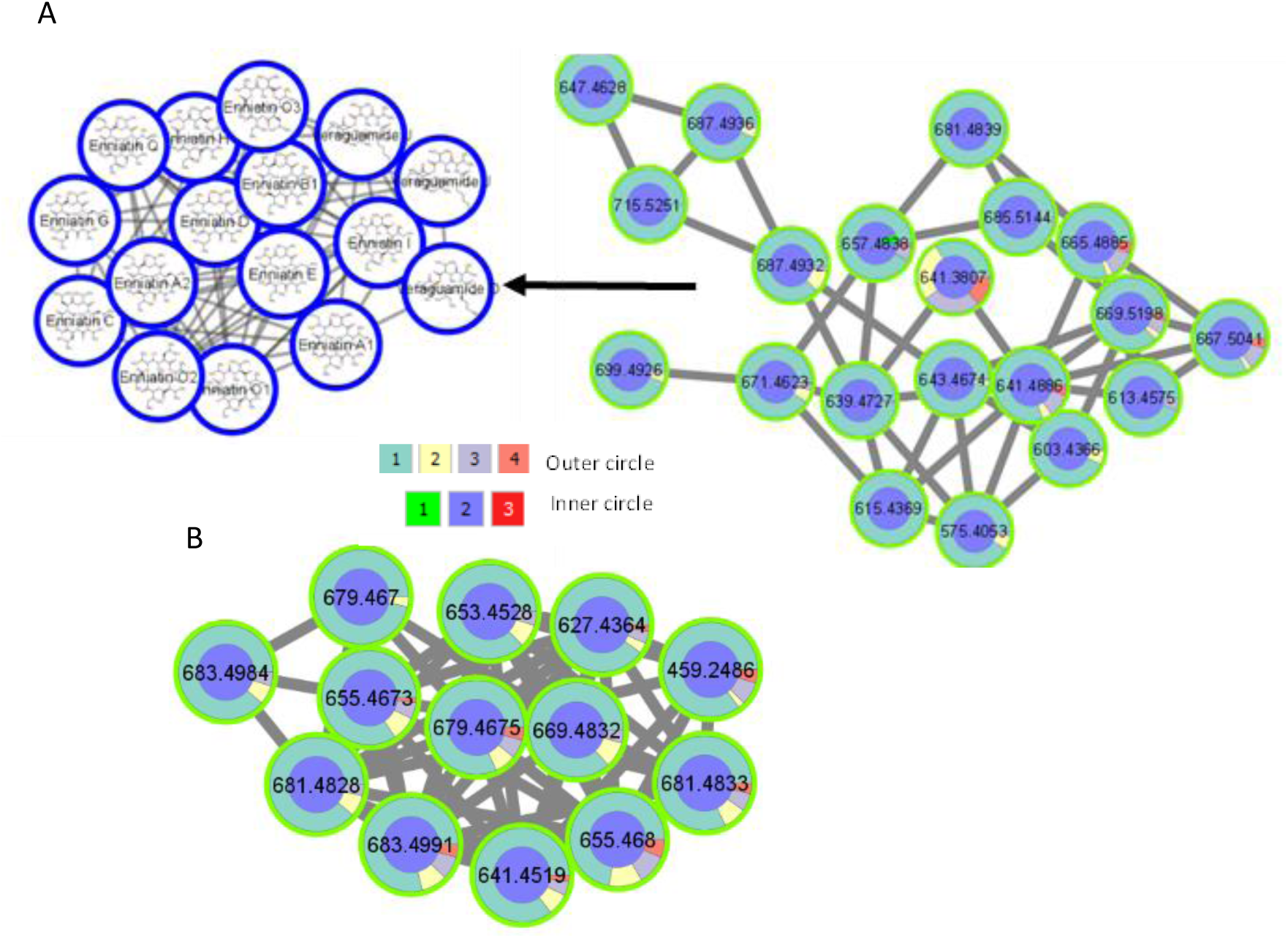
Molecular families from FBMN analysis whose components appeared mainly in the 0% OMSW group. Colors of the inner circle represent features detected in the mushroom substrate (MS), fruiting body (FB) and spent mushroom substrate (SMS) – 1, 2 and 3, respectively. Colors of the outer circle represent features detected in the groups of 0, 33, 60 and 80% OMSW – 1, 2, 3 and 4, respectively. (A) Molecular family annotated by SNAP-MS as enniatins. (B) Molecular family whose components were not annotated by SNAP-MS.

### Validation and fine-tuning of several mushroom metabolite annotations using ^1^HNMR spectra

To verify the in-silico annotations of the here employed computational metabolomics strategies and to potentially resolve isomers, we also performed ^1^H NMR analysis. FB of *H. erinaceus* cultivated on substrate containing 80% OMSW were extracted with methanol, and the crude extract was separated using direct chromatography to obtain eight fractions. The fractions analyzed by ^1^H NMR, and the spectra of three fractions were validated compared to the literature.

Firstly, Fraction 1 included one main chromatogram HPLC peak at RT of 31.35 min (Fig. S7A) with the masses, 557.4203 (M+H) and 579.4020 (M+Na). Briefly summarizing, the masses under the peak were not annotated by GNPS libraries, DEREPLICATOR+, nor NAP. However, using SNAP-MS the peak was annotated as Hericene A (Fig. 6A). Indeed, following fractionation and NMR measurements, the ^1^H-1D-NMR spectrum of the fraction is in line with that of Hericene A (fig. S8),^52^ thereby validating the in-silico annotation.

Secondly, fraction 2 included three chromatogram HPLC peaks at RT of 20.14, 22,125 (main peak) and 25.008 min (Figure S7B) with masses of 595.3996, 571.3991 and 599.4312 respectively for (M+H) and masses of 617.3809, 593.3815 and 621.4122 respectively for (M+Na). The masses under the peaks were not annotated by GNPS libraries nor NAP; however, DEREPLICATOR+ annotated them with Hericenone H, Hericenone F, and Hericenone G, for the peaks appeared at 20.14, 22,125, and 25.008 min., respectively (Fig. 6A). Furthermore, SNAP-MS annotated the masses to be Hericenone E or H for 20.14, Hericenone C or F for peak 22,125 and Hericenone D or G for peak 25.00 ((Fig. S12). Upon comparison of the experimental ^1^H-1D-NMR data (Fig. S9) with literature ^1^H-1D-NMR data, fraction 2 contains Hericenones C, D, and E and not F, G, and H (Figs. S9&S13),^53^ thereby fine-tuning the in-silico annotations by resolving isomeric structures. We refer to Supplementary Text 1 in the Supporting Information for further details.

Finally, fraction 3 included one main HPLC peak at RT of 13.658 (Fig. S7) with masses of 331.1539 for (M+H) and 353.1356 for (M+Na). The masses under the peak were not annotated by GNPS libraries, DEREPLICATOR+, nor NAP; however, SNAP-MS annotated it as Hericenone A (Fig. S11). Comparing the experimental ^1^H NMR data to that of the literature, fraction 3 indeed contains Hericenone A.^54,55^ We do note that the ^1^H NMR spectrum is not clean; however, it contains all the NMR peak shifts related to Hericenone A (Fig. S10), thereby confirming its in-silico annotation.

### Final discussion and conclusions

Computational metabolomics strategies were used for LC-MS/MS data processing and handling to analyze the effect of mixing mushroom substrates with OMSW on *H. erinaceus* and *P. eryngii* specialized metabolic diversity. *H. erinaceus* FB and mycelium were more enriched with specialized secondary metabolites than *P. eryngii* FB and mycelium. FERMO and FBMN results showed that OMSW increased the content of specialized secondary metabolites belonging to the hericenone and erinacerin families in *H. erinaceus* FB and mycelium, respectively. These specialized metabolites have been reported to support health due to their anti-inflammatory, anticancer and neuroprotective properties. The addition of OMSW was further found to decrease the levels of toxic metabolites related to the enniatin family. Altogether, we demonstrate how untargeted metabolomics measurements in combination with computational metabolomics strategies offer a versatile approach to prioritize and structurally annotate affected specialized metabolite families. Given the complexity of most specialized microbial and plant metabolomes, up-to-date mass spectral libraries together with state-of-the-art computational metabolomics strategies are unlikely to cover all metabolites in the natural extracts. However, our analysis demonstrates an innovative approach to better understand their chemical complexities and prioritize relevant metabolite features based on the research question at hand. We do note that the use of various complementary tools can be cumbersome due to their various input demands and parameter settings; however, their complementary perspectives do provide additive value. Furthermore, platforms like FERMO and MolNetEnhancer together with Cytoscape bring together analyses and annotation results, also streamlining the data integration aspects. To further enhance the reliability of in-silico metabolite annotation strategies, NMR validation remains a crucial step, providing high-confidence structural confirmation and complementing MS-based annotations. Incorporating NMR spectroscopy can confirm and fine-tune in-silico annotations, i.e., by resolving isomeric structures, thereby reducing mis-annotations, as we demonstrate for the Hericenone analogues. Hence, NMR analyses nicely complement mass spectrometry-based computational metabolomics workflows. Altogether, the computational metabolomics strategies allowed for an in-depth study of the mushroom-specialized metabolome and the impact of substrate additives thereon, revealing an increase in beneficial specialized metabolites such as hericenone derivatives and a decrease in harmful ones such as enniatins. In conclusion, our study highlights the need for careful consideration of the impact of new nutritional resources, be it beneficial, neutral, or harmful, on the metabolic content of mushrooms or other edible biological products. Furthermore, we provide a framework of computational metabolomics strategies that can be used to assess and compare biochemical diversity of biological products treated in different ways.

## Supporting information

Mzmine batch parameters

## Acknowledgements

The authors thank Niek F. de Jonge (Bioinformatics Group, Wageningen University & Research, Wageningen, the Netherlands), for MS2Query development and integrating it into the FERMO dashboard version 0.8.8. The authors also express their gratitude to the developers of the community-based open-source metabolomics tools that were used in our study.

## Authors Contributions

I.P. and N.E. grow the mushrooms; E.K. extracted the metabolites; R.S. performed the LC-MS/MS measurements; S.K. performed the data and computational metabolomics analysis; M.Z. developed FERMO platform; J.J.J.vdH supervised the research; S.K. wrote the first draft of the manuscript; S.K., I.P. and J.J.J.vdH writing—review and editing. All authors have read and agreed to the final version of the manuscript.

