## Supplemental Figures S1-S13 and Supplementary Text 1 for "Olive mill solid waste induces beneficial mushroom-specialized metabolite diversity revealed by computational metabolomics strategies"

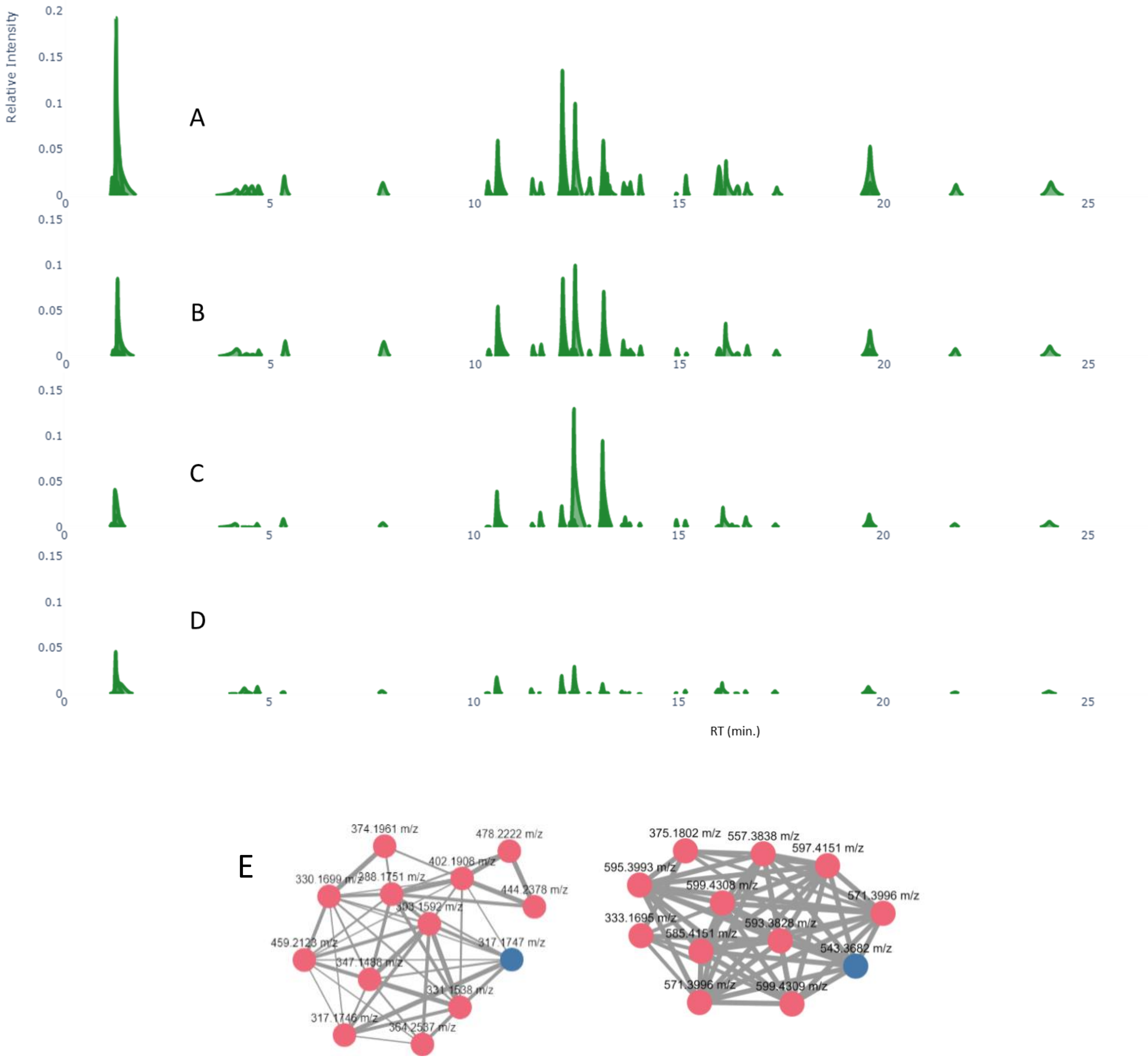


**Figure S1:** FERMO analysis of mass Spectrometry Data from *Hericium* mushroom grown on different percentage of OMSW. Molecular features selected in the FB with quantity higher in the OMSW groups compared to the zero percent OMSW group A) representative sample from the 80 percent OMSW group B) 60 percent OMSW group C) 33 percent OMSW group D) zero percent OMSW group. E) Cytoscape spectral similarity networking of some of the molecular features obtained by FERMO dashboard.

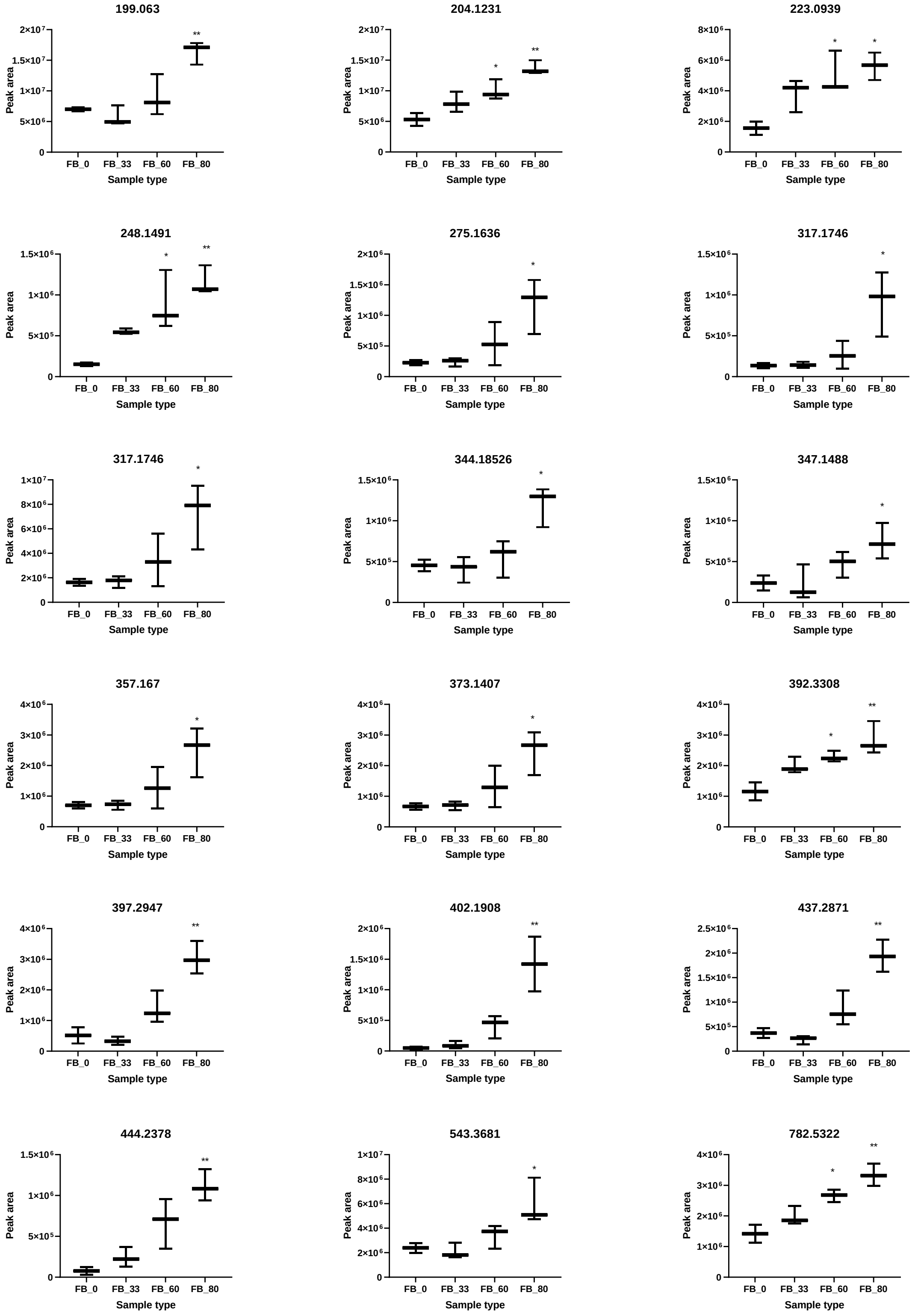


**Figure S2:** Selected metabolite features which are significantly increased in Hericium fruiting body grown on mushroom substrate mixed with 0, 33, 60 and 80 percent of OMSW (FB_0, FB_33, FB_60 and FB_80 respectively). The figures present the m/z on top, and mean area values ± SD from three extracts for each group ($n$ = 3). *$P \leq 0.05$, **$P \leq 0.01$ in relative to the FB_0 group. Statistical analysis was by one-way ANOVA and Graphpad prism 9 software.

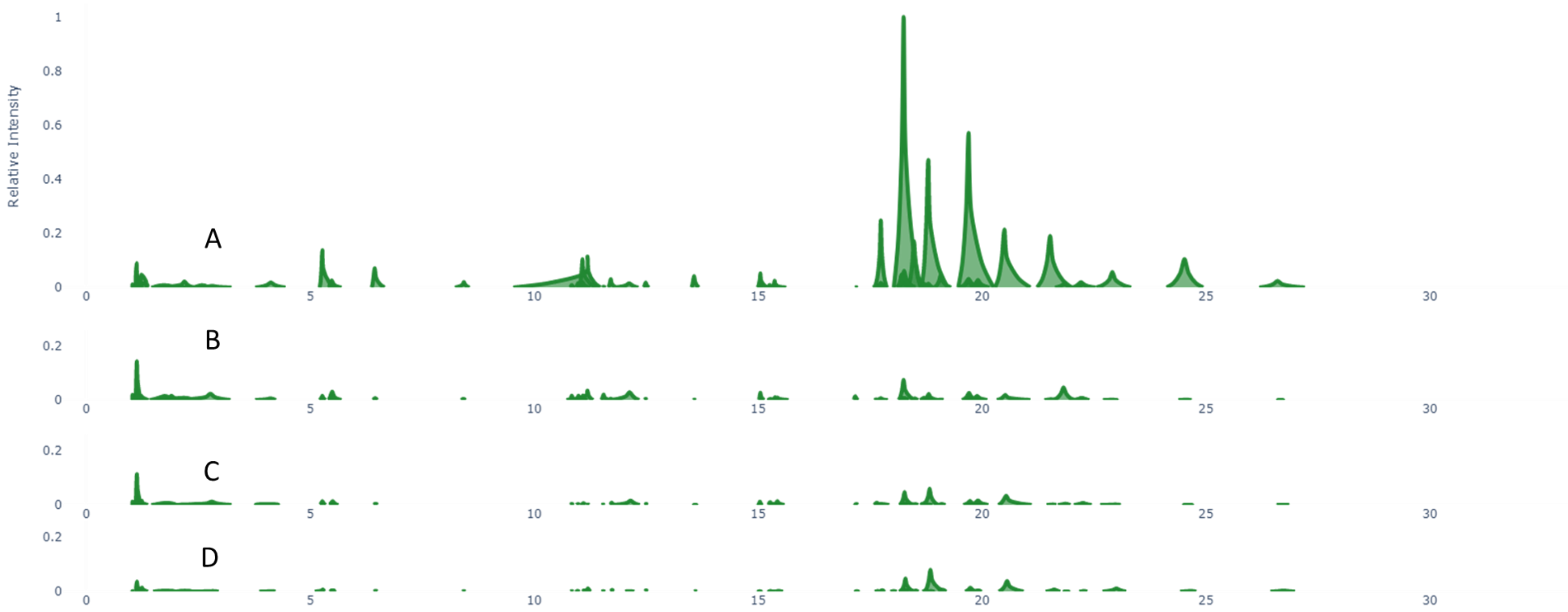


**Figure S3:** FERMO analysis of mass spectrometry profiles from Hericium mushroom fruiting body grown on different percentage of OMSW. Selected molecular features being at least four-fold higher in the zero percent OMSW group compared to the 80 percent OMSW group displayed in representative samples from the A) zero percent OMSW B) 33 percent OMSW C) 60 percent OMSW D) 80 percent OMSW.

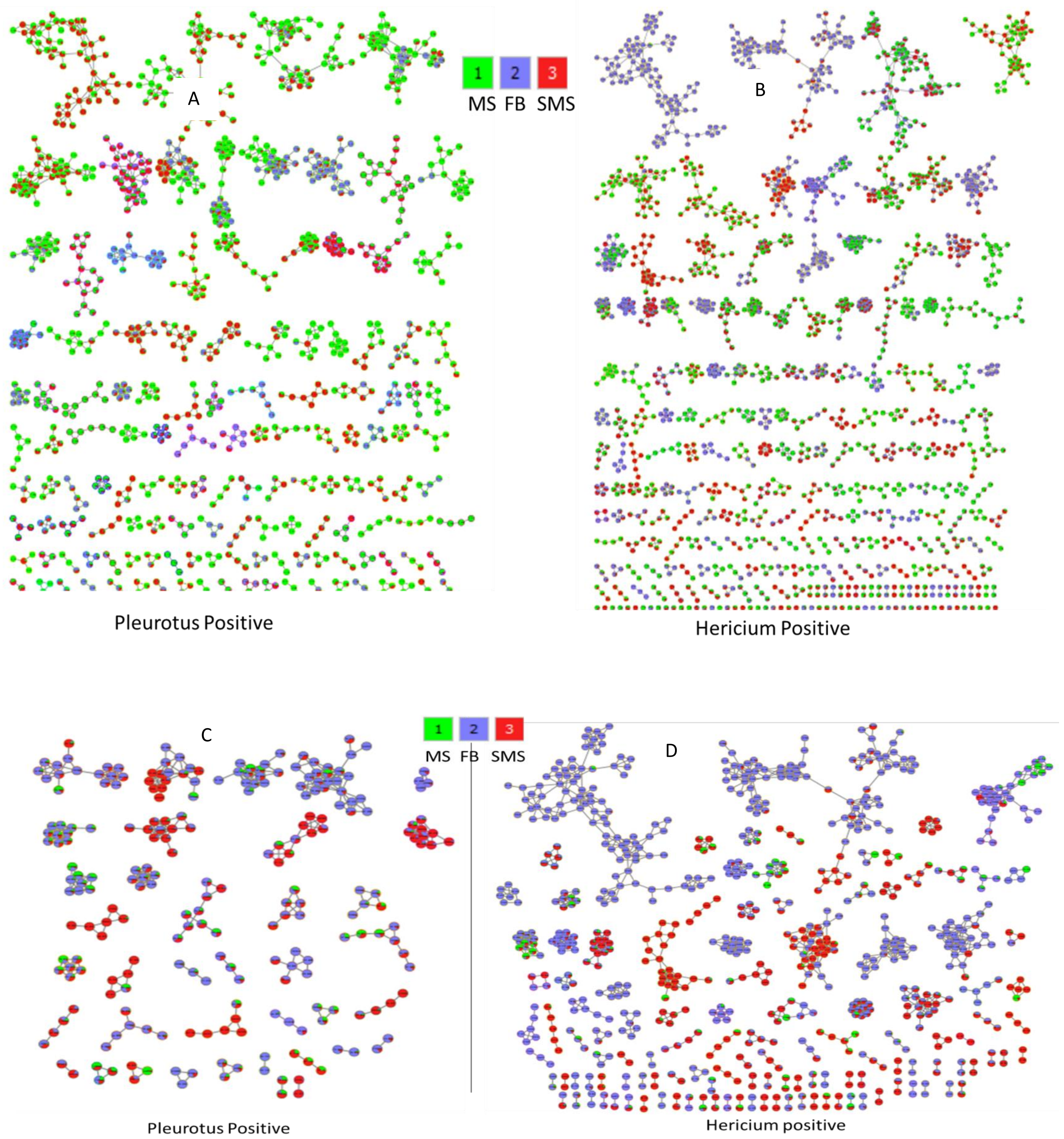


**Figure S4:** Features based molecular networking (FBMN) for mushroom fruit bodies (FB), spent mushroom substrates (SMS) and mushroom substrates (MS) for Pleurotus (A) and Hericium (B). (C) and (D) are FBMN from fruit body (FB) and spent mushroom substrate (SMS) after removing molecular networks from the mushroom substrate (MS) for Pleurotus and Hericium respectively.

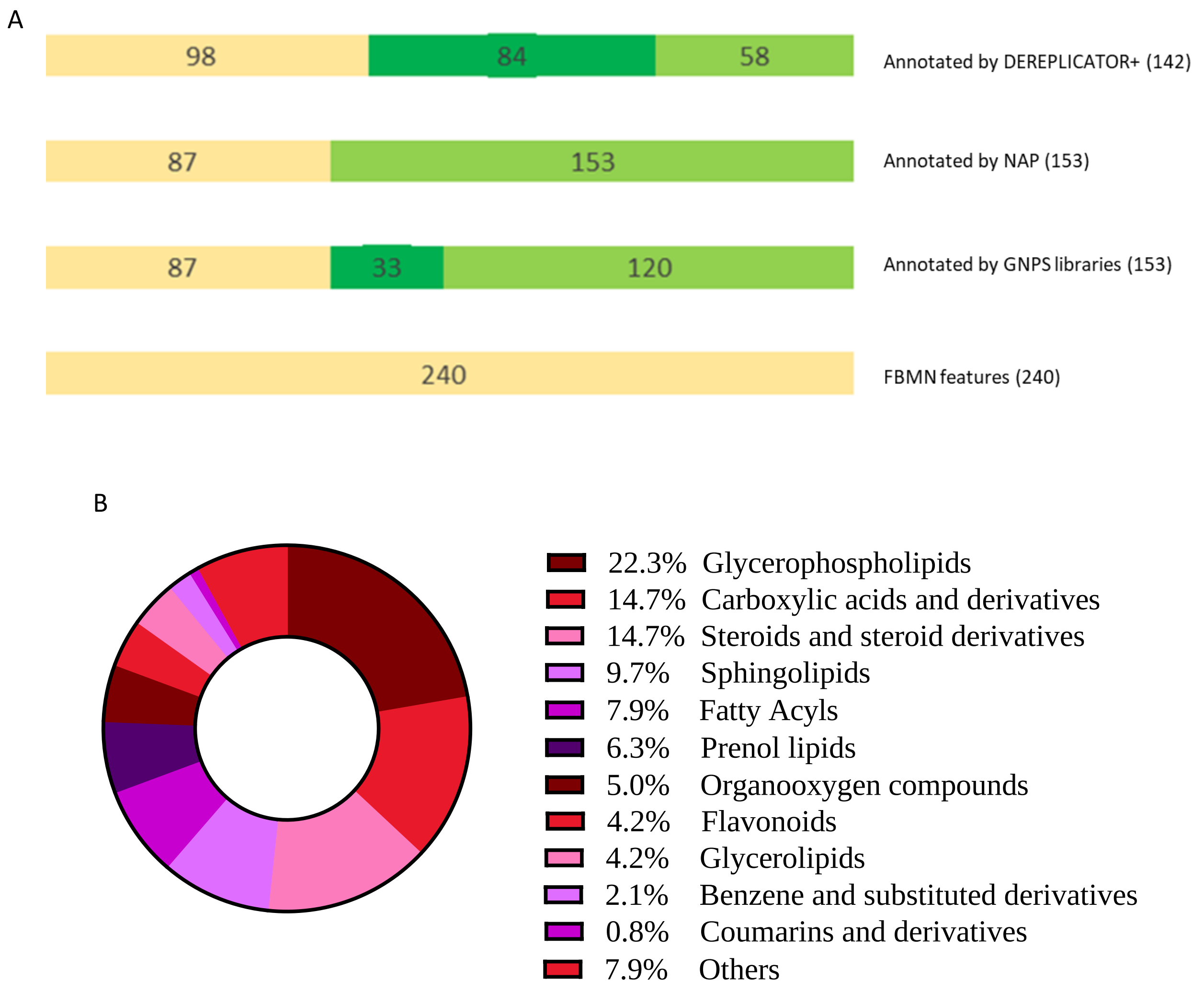


**Fig. S5.** (A) Annotation results obtained from Pleurotus FBMN analysis using GNPS libraries, DEREPLICATOR+ and NAP. The yellow color represents the number of detected features, the light green color the annotated features, and the dark green color the annotated features with score higher than 0.95 for GNPS libraries and higher than 12 for DEREPLECATOR plus database (B) Chemical classification (using ClassyFire) of the molecular families obtained from the Pleurotus FBMN.

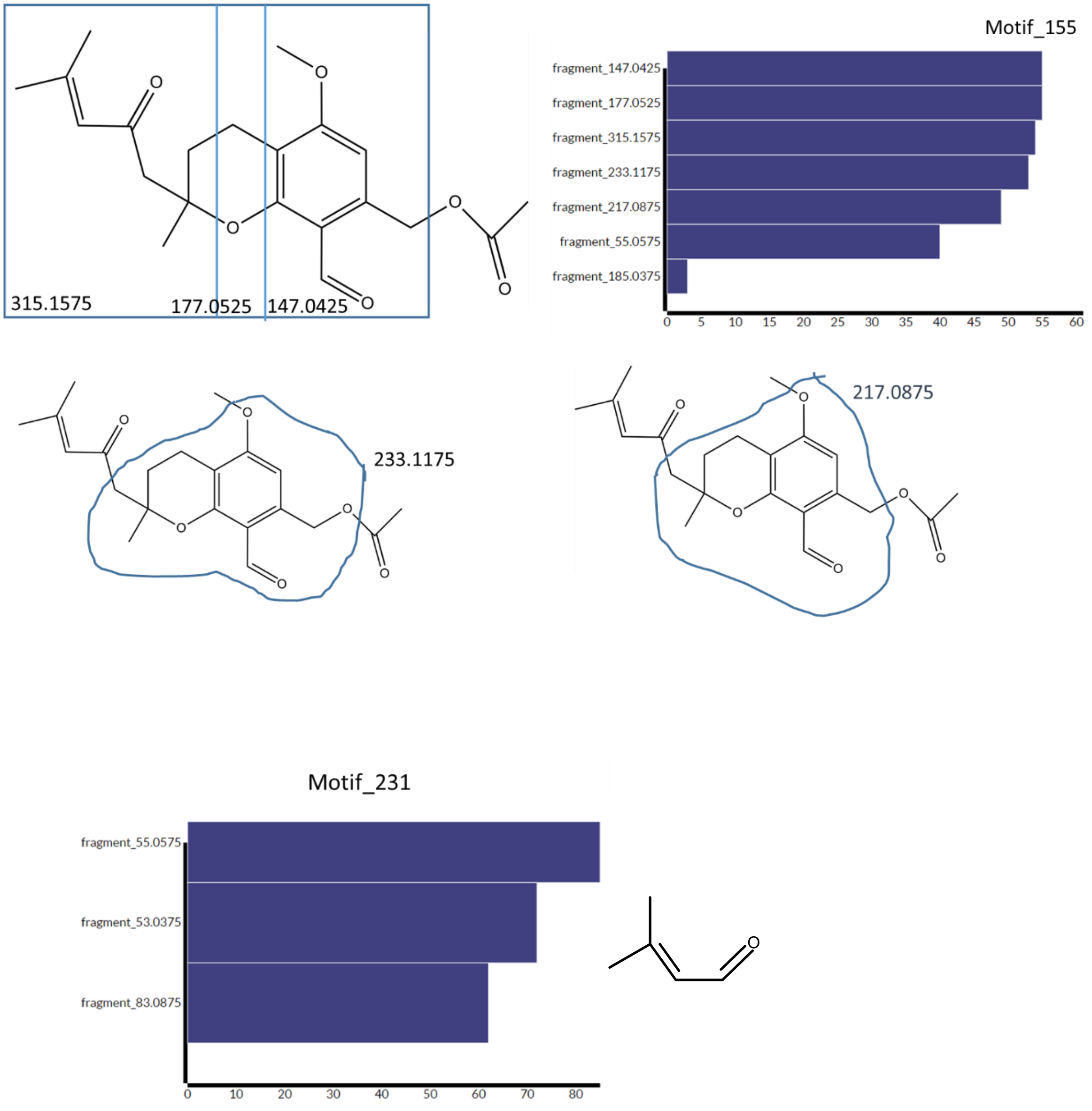


**Figure S6:** MS2LDA discovered motifs that were detected in most of the nodes in the molecular networking which annotated as analogs of hericenones.

A

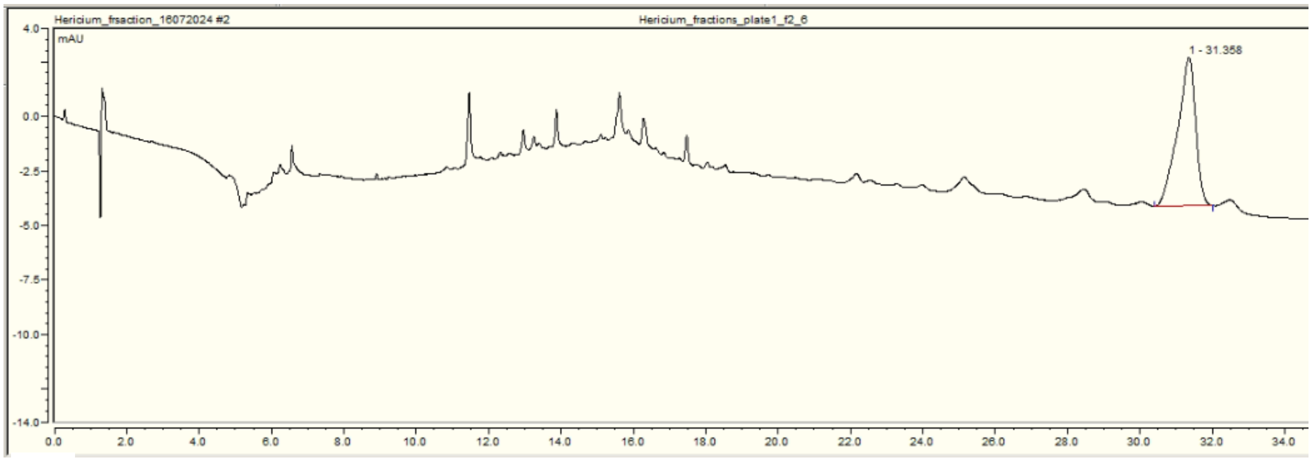


B

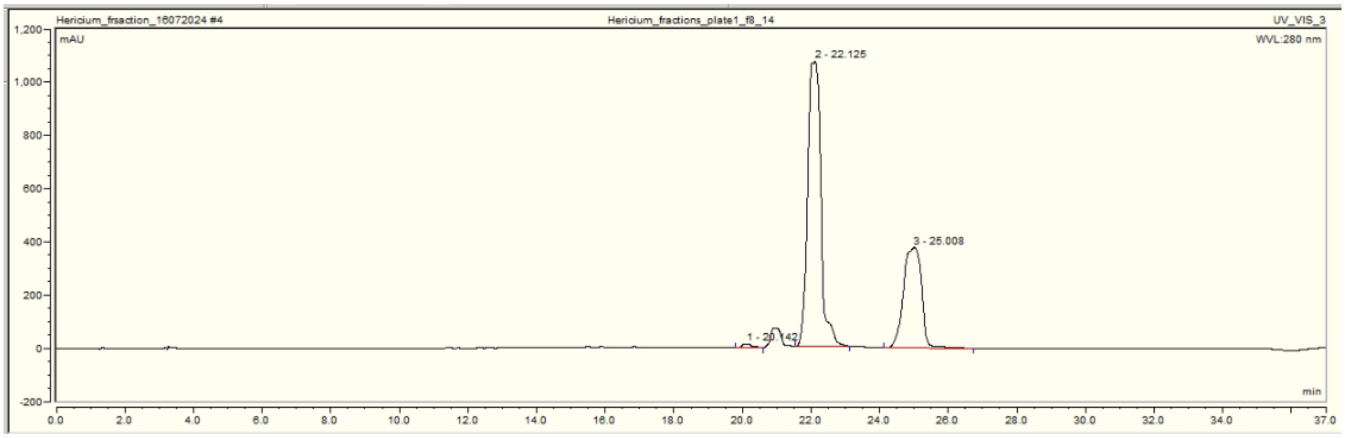


C

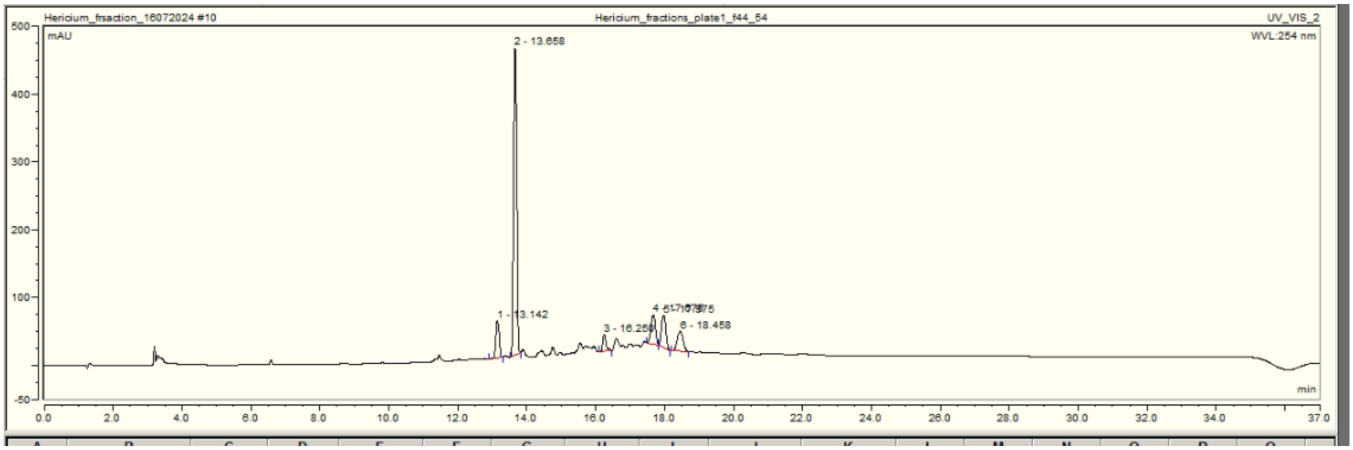


**Figure S7:** HPLC chromatograms of fractions 1 (A), 2(B), and 3(C)

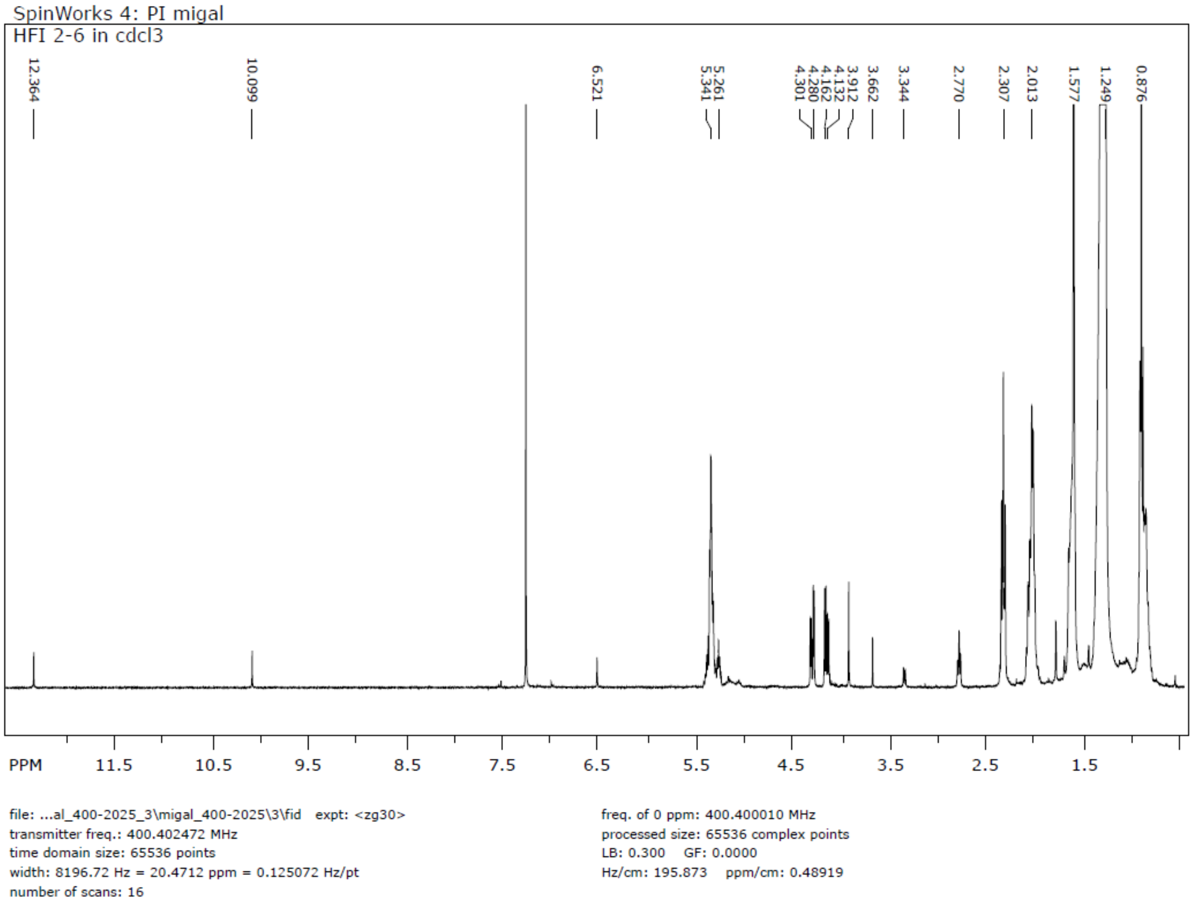


**Figure S8:** NMR spectrum of fraction 1.

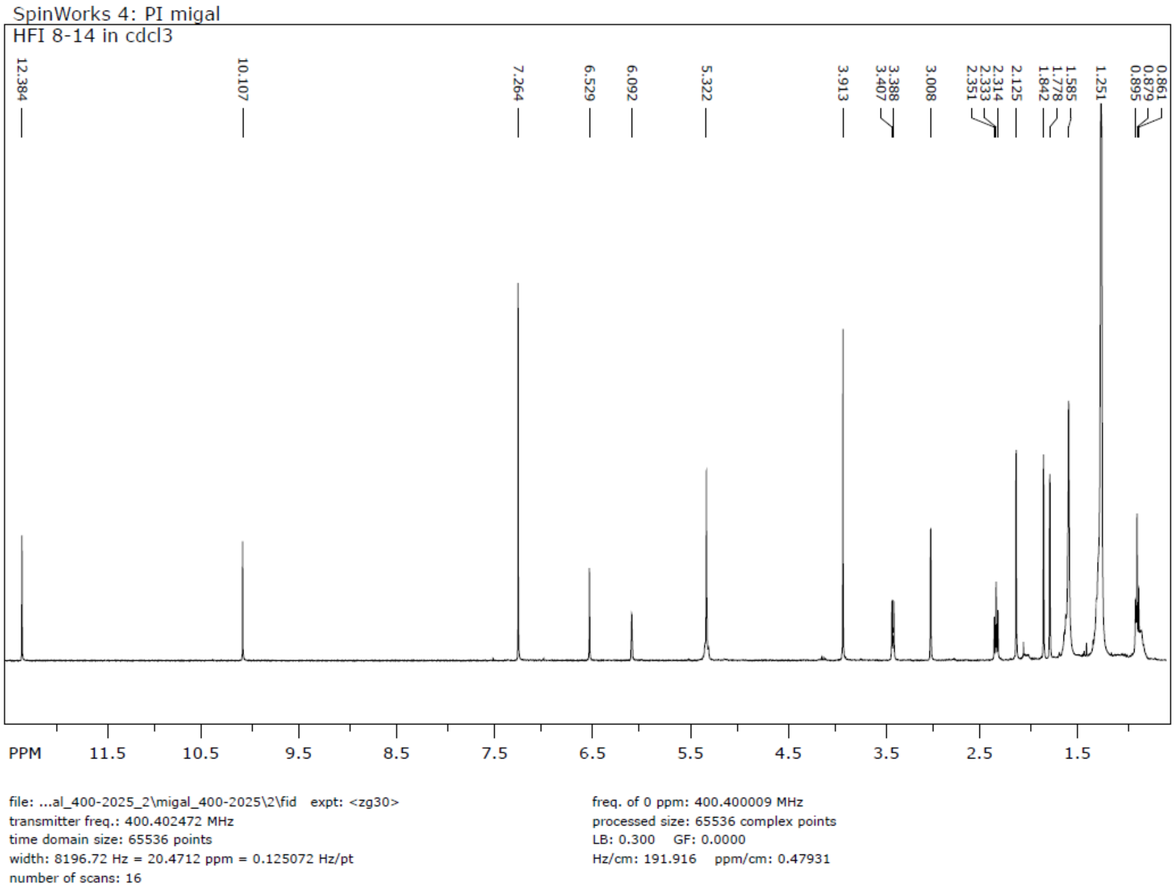


**Figure S9:** NMR spectrum of fraction 2.

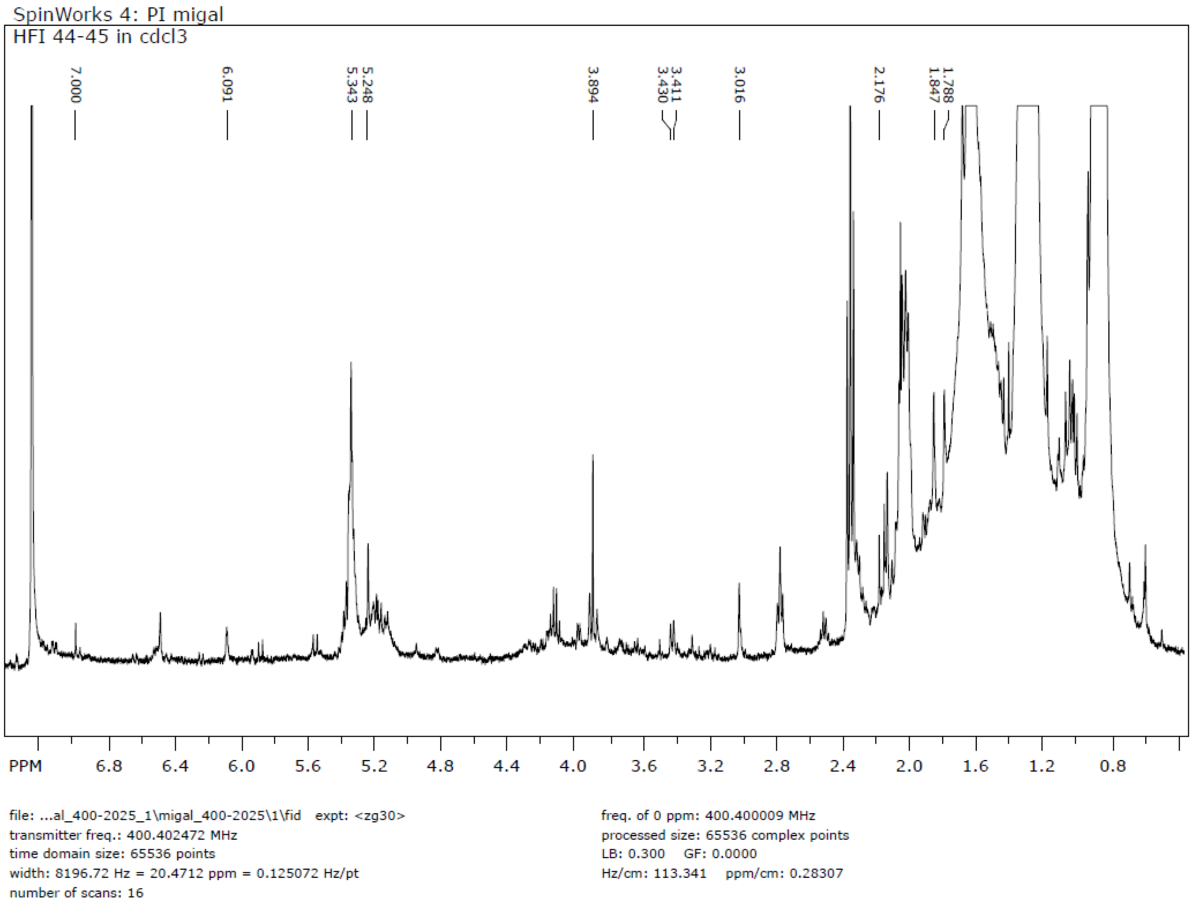


**Figure S10:** NMR spectrum of fraction 3.

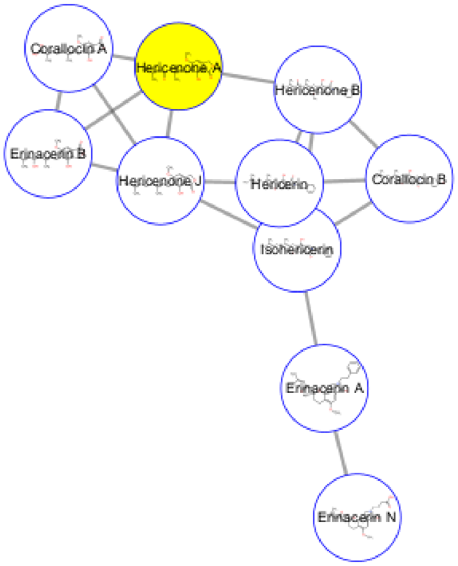


**Figure S11:** Molecular network obtained from SNAP-MS includes annotation of the mass 331.1539 to be Hericenone A.

### Supplementary Text 1 – Hericenone isomer annotations and validation using $^1$H NMR

Fraction 2 includes three HPLC chromatographic peaks at RT of 20.14, 22,125 (main peak) and 25.008 min (Figure S7B) with masses of 595.3996, 571.3991 and 599.4312 respectively for (M+H) and masses of 617.3809, 593.3815 and 621.4122 respectively for (M+Na). Briefly summarizing, the masses under the peaks were not annotated by GNPS libraries, and NAP. They annotated by DEREPLICATOR+ to be Hericenone H, Hericenone F, and Hericenone G for the peaks appeared at 20.14, 22,125, and 25.008 min, respectively. Using SNAP-MS it was annotated to be Hericenone E or H for 20.14, Hericenone C or F for peak 22,125 and Hericenone D or G for peak 25.00 (Fig. S12 A, B and C).

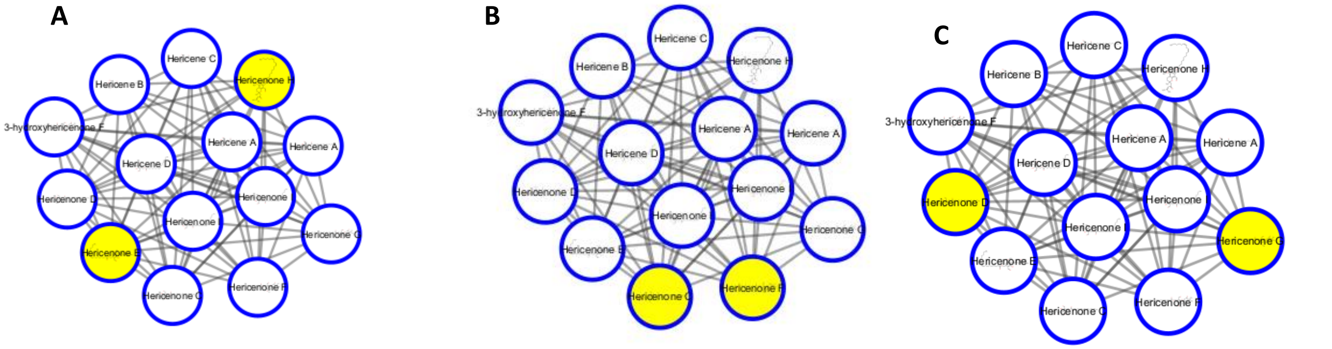


**Figure S12**: Annotation using SNAP-MS for the peaks appeared at 20.14 (A), 22,125 (B)and 25.008 min (C).

When comparing to literature values, we found that the $^{1}$H-1D-NMR spectrum of fraction 2 (Fig. S9) contained peak shifts in line with those of Hericenones C, D, and E, and not with those of F, G, and H (Fig. S13). For example, Hericenones C, D, and E includes the peak shift of 12.3 and 10.1 and others while Hericenones F, G and H contains the peak shift at 10.4 ppm.  Fraction 2 spectrum is similar to that of Hericenones C, D, and E, and it does not contain the peak at 10.4 ppm (Fig. S9 and S13).

Because of the similarity between the $^{1}$H-1D-NMR spectrum of the fraction and $^{1}$H-1D-NMR spectrum of Hericenones C, D and E, we can conclude that the chromatographic peak at 20.14 with the masses 595.3996 and 617.3809 can be annotated with Hericenone E, the chromatographic peak at 22,125 with the masses 571.3991 and 593.3815 as Hericenone C, and the chromatographic peak 25.008 with the masses 599.4312 and 621.4122 as Hericenone E.

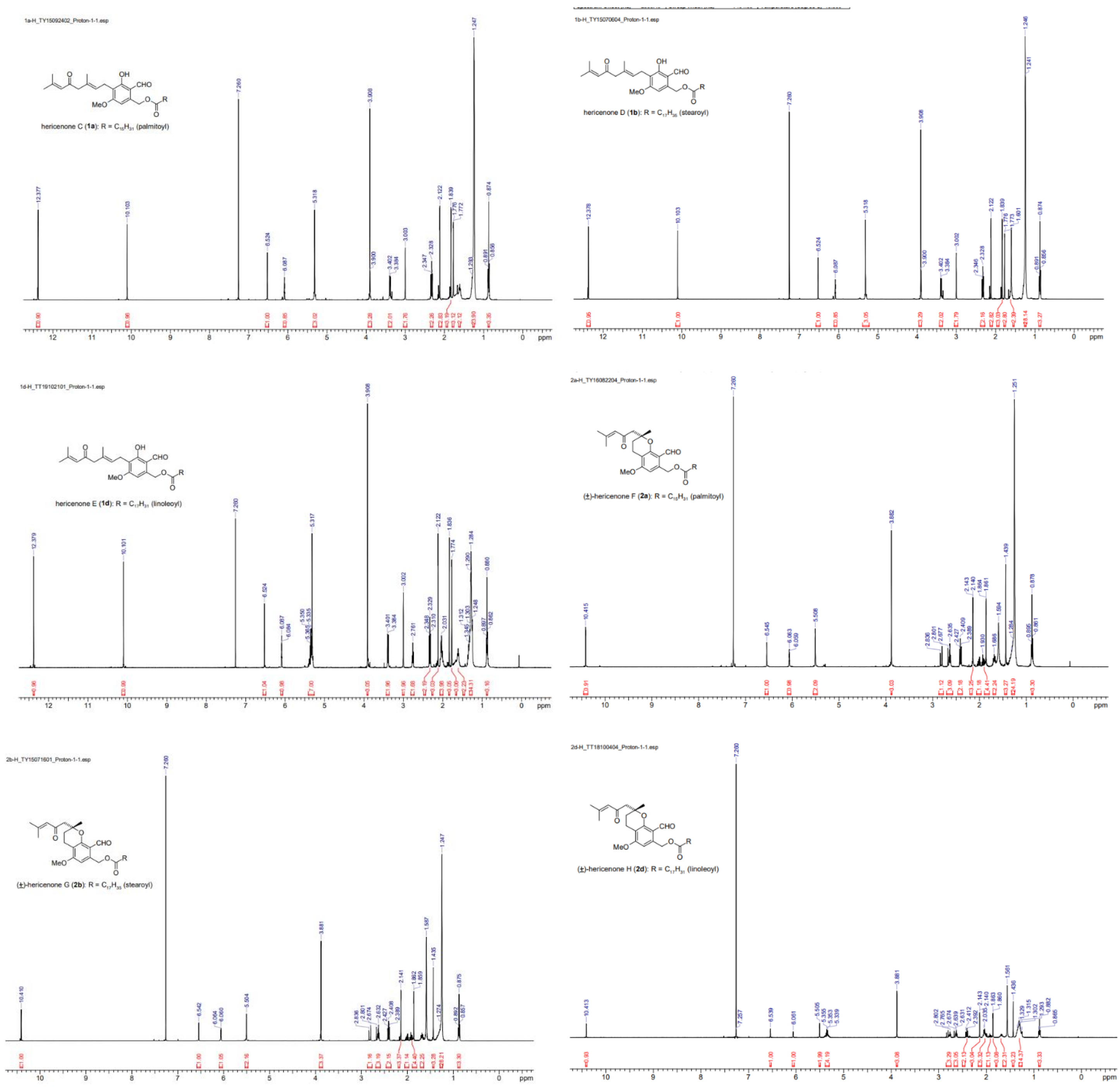


**Figure S13**: NMR spectrum of Hericenones C, D, E, F, G, H from the literature (Kobayashi S, Tamura T, Koshishiba M, et al. Total Synthesis, Structure Revision, and Neuroprotective Effect of Hericenones C-H and Their Derivatives. J Org Chem 2021;86:2602-20.).
